# From Pixels to Phenotypes: Integrating Image-Based Profiling with Cell Health Data Improves Interpretability

**DOI:** 10.1101/2023.07.14.549031

**Authors:** Srijit Seal, Jordi Carreras-Puigvert, Anne E Carpenter, Ola Spjuth, Andreas Bender

**Affiliations:** Yusuf Hamied Department of Chemistry, University of Cambridge, Cambridge, United Kingdom; Department of Pharmaceutical Biosciences and Science for Life Laboratory, Uppsala University, Uppsala, Sweden; Imaging Platform, Broad Institute of MIT and Harvard, Cambridge MA, USA

**Keywords:** Cell Painting, Cell Health, BioMorph, Toxicity Prediction, Cell Morphology, Interpretability, Image-based profiling, machine learning

## Abstract

Cell Painting assays generate morphological profiles that are versatile descriptors of biological systems and have been used to predict *in vitro* and *in vivo* drug effects. However, Cell Painting features are based on image statistics, and are, therefore, often not readily biologically interpretable. In this study, we introduce an approach that maps specific Cell Painting features into the BioMorph space using readouts from comprehensive Cell Health assays. We validated that the resulting BioMorph space effectively connected compounds not only with the morphological features associated with their bioactivity but with deeper insights into phenotypic characteristics and cellular processes associated with the given bioactivity. The BioMorph space revealed the mechanism of action for individual compounds, including dual-acting compounds such as emetine, an inhibitor of both protein synthesis and DNA replication. In summary, BioMorph space offers a more biologically relevant way to interpret cell morphological features from the Cell Painting assays and to generate hypotheses for experimental validation.

**GRAPHICAL ABSTRACT:** 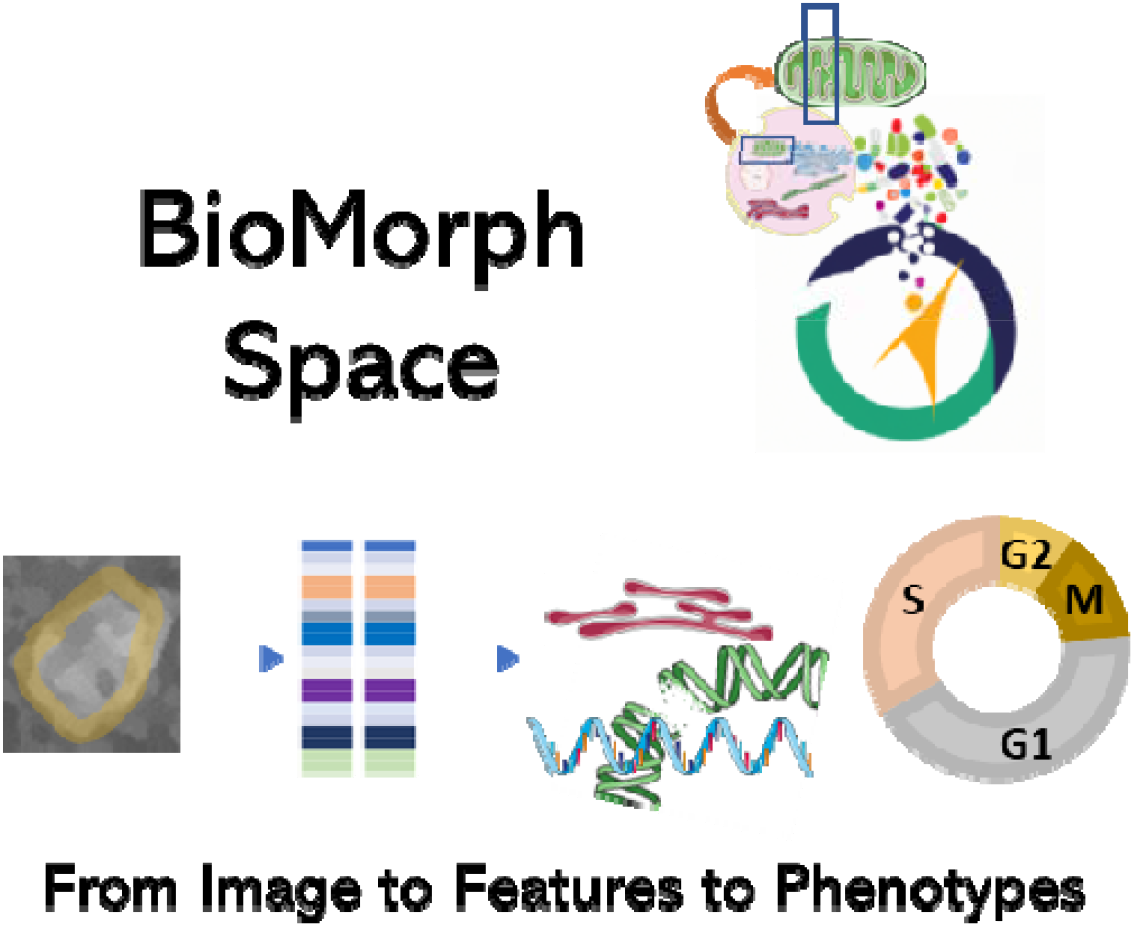

**IN BRIEF:** Seal et al. used machine learning models and feature selection approaches to group cell morphological features from Cell Painting assays and to describe the shared role of these morphological features in various cell health phenotypes. The resulting BioMorph space improves the ability to understand the mechanism of action and toxicity of compounds and to generate hypotheses to guide future experiments.

**HIGHLIGHTS:**

- Combining Cell Painting and Cell Health imaging data defines the BioMorph space.
- BioMorph space allows detecting less common mechanisms for bioactive compounds.
- BioMorph space can generate MOA hypotheses to guide experimental validation.
- BioMorph space is more biologically relevant and interpretable than Cell Painting features.

## INTRODUCTION

Cell Painting profiles^1^ can be used to study the morphological characteristics of cells treated with chemical or genetic perturbations and provide valuable information about the function of a biological system.^2,3^ The Cell Painting assay involves labelling eight relevant cellular components or organelles with six fluorescent dyes, imaging them in five channels,^4^ and analysing images^5^ to provide thousands of morphological features such as shape, area, intensity, texture, correlation, etc. Cell Painting data have been used to successfully predict drug effects on many aspects of cell health^6^, such as cytotoxicity^7^, mitochondrial toxicity^8^, proteolysis targeting chimera (PROTAC) phenotypic signatures^9^, and other types of bioactivities^10,11^. Further, Cell Painting data can be used to cluster together compounds with various mechanisms of action based on the similarity of resulting morphological features they induce.^12,13^ Thus, Cell Painting features serve as a tool for investigating the chemical space and enabling the prediction of a compound’s biological activities.^14,15^

In general, Cell Painting features are obtained using classical image processing software, such as CellProfiler.^5^ After establishing the threshold for distinguishing signal from the background noise, classical image processing software identifies all signal-containing pixels and their intensity, and groups neighbouring pixels into objects using object-based correlations.^16^ The measured morphological features are then extracted from each object (cell or subcellular structure). Given this image processing pipeline, Cell Painting features primarily represent numerical data from image analysis (often aggregated to the treatment level for machine learning tasks), rather than directly reflecting the underlying biological processes or molecular interactions.^17^ Therefore, interpreting the Cell Painting data and making informed decisions about drug safety, toxicity, efficacy, or the underlying mechanisms and cellular processes based on such data remains challenging. This suggests that integrating Cell Painting features with some *a priori* knowledge about the biological effects of different chemical or genetic perturbations may result in improved predictive power of models derived from Cell Painting data.

An orthogonal strategy that considers *a priori* knowledge about the biological effects is the Cell Health assay, a set of two image-based assays^18^ that collectively capture a broad range of biological pathways. The Cell Health assay thus records measurable characteristics from cellular responses to different treatments (or environmental conditions, pathological states etc.)^19,20^ which determine the overall condition, functionality, and viability of cells^21^, including the different stages of the cell cycle. Following a similar premise, a study by Way et al used the Cell Health assay and CRISPR/Cas9 to genetically perturb a small subset of 118 gene perturbations across three cell lines.^6^ Recording the effects of these genetic perturbations using carefully chosen reagents for specific cellular processes (*e.g.* apoptosis, DNA damage, etc.) allowed them to define 70 Cell Health readouts that can be used to quantify and model cellular responses to different treatments.^6^ The Cell Health readouts are directly related to mechanisms and cellular function and can be used to predict the mechanism of action (MOA) of the perturbation and derive functional conclusions. However, unlike the hypothesis-free Cell Painting assay, the Cell Health assay requires specifically targeted reagents focused on individual measurement and is difficult to scale for high throughput applications.

Here, we address the limitations of both Cell Painting and Cell Heath assays by integrating their capabilities to define a new BioMorph space that provides a function-informed framework for interpreting Cell Painting features in the cell biology context. We used publicly available Cell Painting data^6^ and Cell Health data^6^ to define this BioMorph space. To demonstrate the use of the BioMorph space, we used the Cell Painting features from chemical perturbations^4^ to predict a range of nine broad biological activities from ToxCast, such as apoptosis, cytotoxicity, oxidative stress, and ER stress. We then mapped important Cell Painting features from these models into BioMorph terms. Identifying the BioMorph terms that contribute most strongly to model performance helped generate MOA hypotheses, some in agreement with the existing literature and some novel. Taken together, our proposed method offers several potential advantages, including improved interpretability of cell morphology features, enhanced understanding of cellular mechanisms and MOA, and more interpretable predictions of drug toxicity and efficacy.

## RESULTS AND DISCUSSION

We developed a structured framework for mapping Cell Painting features to a more biologically synthesized BioMorph space. We used feature selection, linear regression, and Random Forest classifiers on the publicly available Cell Painting and Cell Health datasets^6^ for a set of 119 CRISPR perturbations (for further details see Methods). This mapping was then used to interpret models predicting biological activity using a dataset containing morphological profiles of 30,000 small molecules produced using the Cell Painting assay^22^. Mapping those Cell Painting features that contribute the most to the performance to the BioMorph space led to an improvement in interpretability and allowed us to generate hypotheses on the cause of cellular effects.

### Development of the BioMorph space through the integration of Cell Painting and Cell Health assays

We mapped the groups of Cell Painting features into five levels within the BioMorph space as shown in Figure 1 (see Methods for technical details and Supplementary Table S1 and Figure S1 for all terms). These levels were chosen to leverage the maximum information from the Cell Health assay and include the Cell Health assay type (Level 1), Cell Health measurement type (Level 2), specific Cell Health phenotypes (Level 3), Cell process affected (Level 4), and the subset of Cell Painting features (Level 5). The first level, the Cell Health assay type, represents results from one of the two screening assays used to measure the Cell Health parameters, *e.g.* the viability assay or the cell cycle assay. The second level, Cell Health measurement type, describes the various aspects of Cell Health measured in that assay, such as cell death, apoptosis, reactive oxygen species (ROS), and shape for viability assays, and cell viability, DNA damage, S phase, G_1_ phase, G_2_ phase, early mitosis, mitosis, late mitosis, and cell cycle count for cell cycle and DNA damage assays. The third level, specific Cell Health phenotypes, describes specific assay readouts that capture different aspects of the phenotype, such as the fraction of cells in G_1_, G_2_ or S-phase cells. The fourth level, the Cell process affected, contains information on the type of Cell process affected that caused the change in morphological characteristics, e.g., effects of chromatin modifier, DNA damage, metabolism, etc. Finally, the fifth level, Cell Painting features, is the subset of Cell Painting image-based features that map to the combination of the previous four levels. These five levels formed the basis of the BioMorph space.

**Figure 1.**
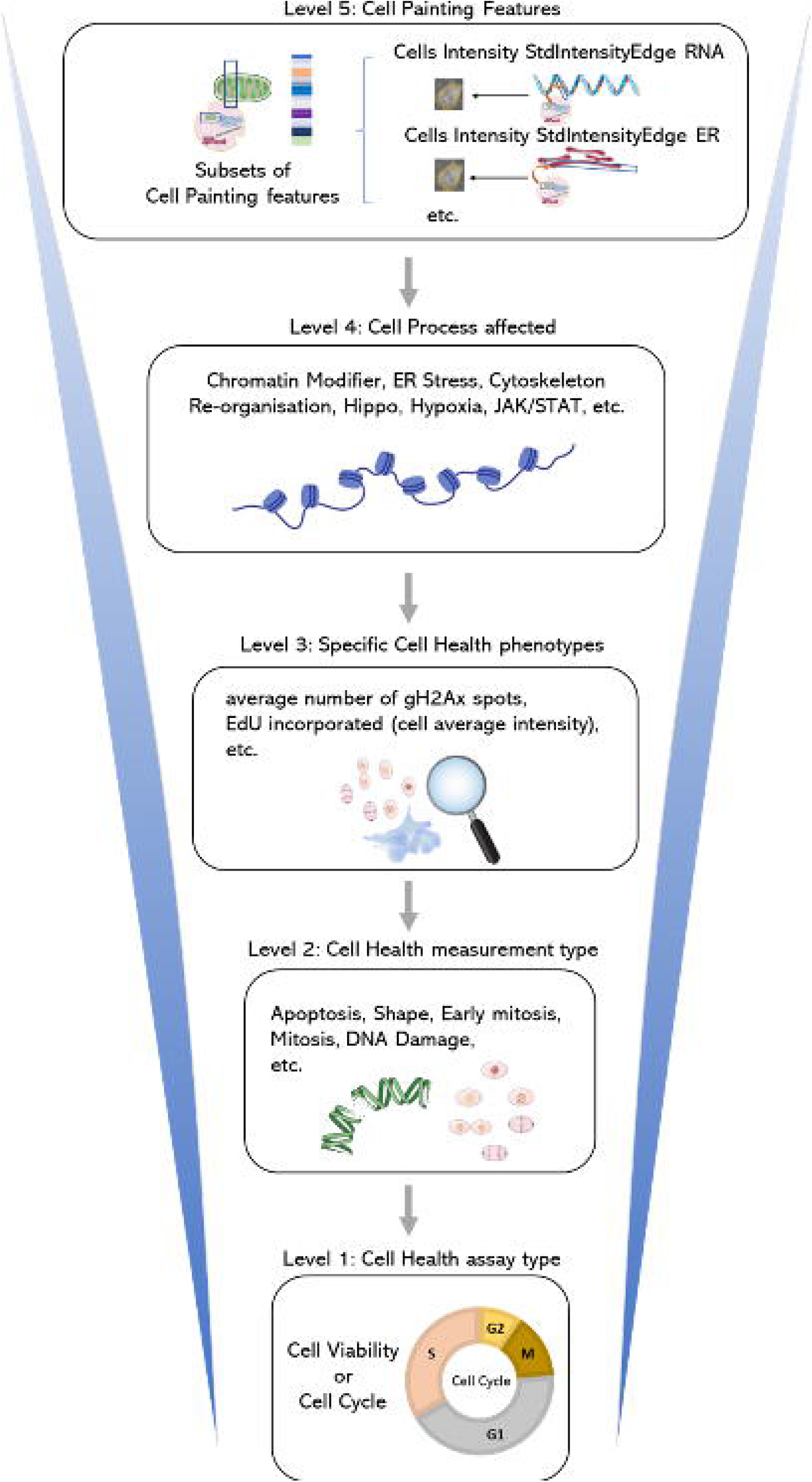
A map of the BioMorph space. A general representation of the hierarchy of levels is shown (with examples) for each BioMorph term that is organised from Cell Painting features (level 5) and containing information on Cell Health (level 3) associated with measurement type (level 2) under an assay type (level 1) associated with Cell process affected (level 4). Further terms in Supplementary Figure S1 with all terms listed in Supplementary Table S1

To build the BioMorph space we focused on the overlap of perturbations between Cell Painting and Cell Health assay containing 827 Cell Painting features and 70 continuous Cell Health endpoints. We used an all-relevant feature selection method Borutapy^23^ (Figure 2, step A) to detect a subset of Cell Painting image features that contain information important for predicting each of the 70 Cell Health labels. Further, we trained a baseline Linear Regression model (Figure 2, step B) and determined which subsets of Cell Paintings features are relatively better predictors for each of the 70 Cell Health labels. Meaningful models were built for 34 Cell Health labels which resulted in corresponding 34 subsets of Cell Painting features. Next, for each of the Cell Health labels, we used Borutapy to select subsets of Cell Painting features that could distinguish a particular CRISPR perturbation from the negative controls (Figure 2, step C). Lastly, we trained a baseline Random Forest Classifier (Figure 2, step D) to predict which these sets of selected Cell Painting features perform better at differentiating negative controls from respective CRISPR perturbations (MCC>0.50). This led to 412 subsets (combinations of the various levels above) of informative Cell Painting features which were used to define 412 BioMorph terms (Figure S1 and Supplementary Table S1 lists all the terms and their description; Figure 2, step E). Thus, each BioMorph term integrates a unique combination of information derived from the perturbations and Cell Health labels in the Cell Health assay and a subset of Cell Painting features.

**Figure 2.**
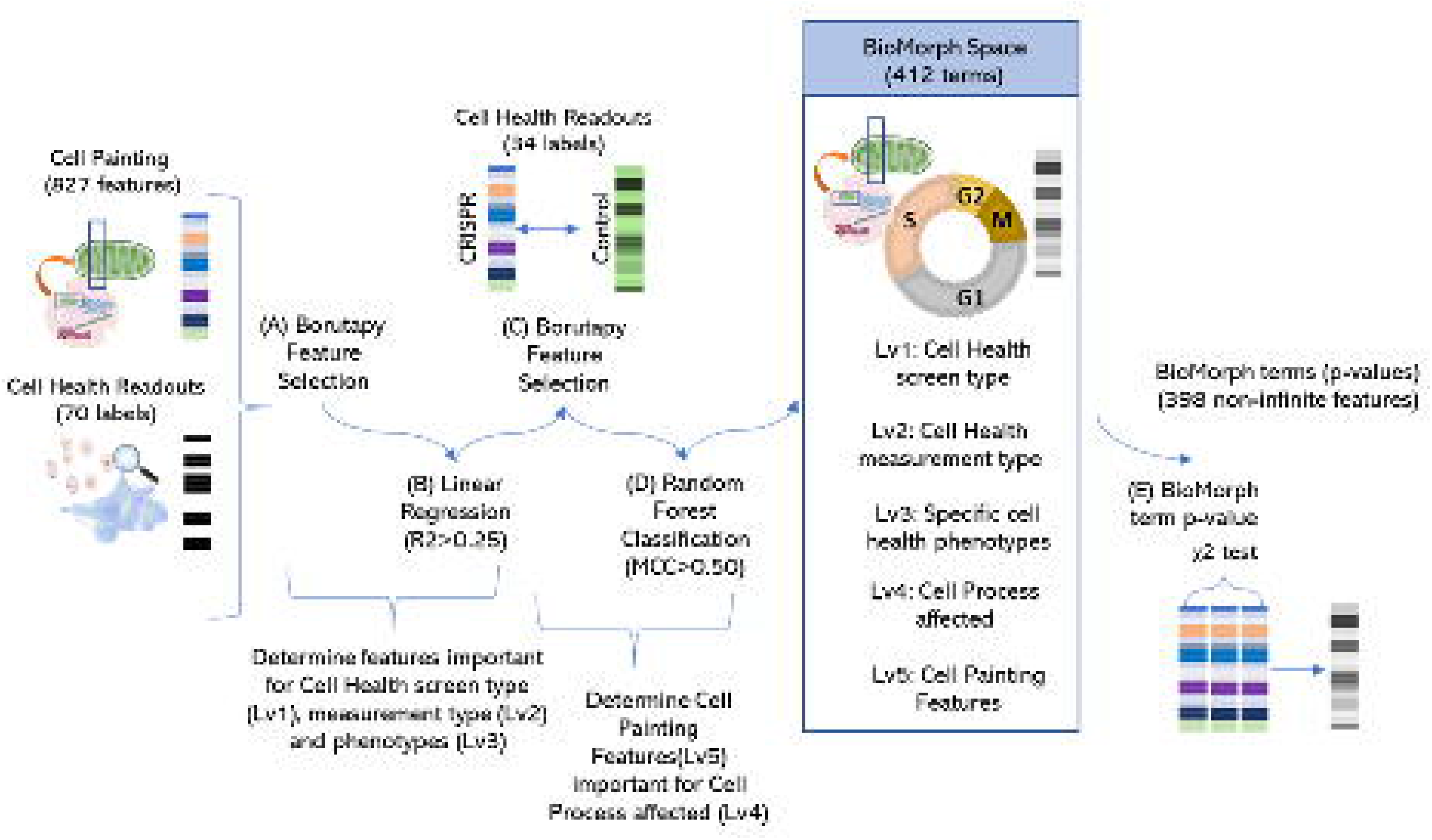
Schematic representation of methodology to generate BioMorph terms mapped from CRISPR perturbations measured by the Cell Painting assay and Cell Health assay. Further details on all terms terms in Supplementary Figure S1 with all BioMorph Space listed in Supplementary Table S1

For example, the BioMorph term “viability_apoptosis_vb_percent_dead_only_Chromatin Modifiers” records a morphological change that includes information about the “fraction of caspase negative in dead cells” (level 3) associated with apoptosis (level 1), measurement type (level 2), and the effect of CRISPR knockout of a gene associated with a chromatin modifier benchmarked against the negative control (level 4) for which a particular set of Cell Painting features (level 5) contained a signal to distinguish from negative control. This multi-level approach allows for a more nuanced understanding of cellular health and its relation to specific biological mechanisms. In the example given above, the caspase-negative dead cells are a readout for cells that have undergone non-apoptotic cell death.^24^ Furthermore, the term associates this form of cell death with the effects of the CRISPR knockout of a gene associated with a chromatin modifier, which is consistent with existing evidence that certain inhibitors that affect chromatin modifications, such as histone deacetylase (HDAC) inhibitors, can initiate non-apoptotic cell death mechanisms.^25^ Therefore, this specific BioMorph term captures signals associated with these biological characteristics and MOA.

### BioMorph space retains all information for biological activity from the original Cell Painting features

We first ensured that BioMorph space contains all information from the original Cell Painting readouts, which we found to be the case as shown in Supplementary Figure S2. We used Random Forest classifiers using 398 BioMorph terms directly as features (p-values from a χ2 test on the groups of Cell Painting features; although there were 412 terms defined, only 398 terms out of these were non-infinite and continuous and used for modelling). We compared these classifiers to the models trained on all 827 Cell Painting features. Supplementary Table S2 shows the mean AUC-ROC and mean balanced accuracy from the 20 internal test sets of the repeated nested cross-validation for all 9 biological activities. Overall, models using Cell Painting features (mean AUC=0.60) achieved a similar performance compared with models using BioMorph terms (mean AUC=0.61) (as shown in Supplementary Figure S2 with a paired t-test). Thus, transforming important Cell Painting features from models into the BioMorph space made these models more interpretable without any loss in performance compared to models using BioMorph terms directly.

### Incorporating information about phenotypic characteristics (Cell Health phenotype; level 3) enhances the ability to connect Cell Painting features (level 5) to biological activity from ToxCast

To compare the ability of Cell Painting features alone, or when integrated with Cell Health phenotypes (level 3), to predict biological activity, we used 56 cytotoxicity and cell stress response assays from a public dataset called ToxCast ^26^. We generated predictions for nine biological activities (see Judson et al^56^ for mapping 56 assays into nine activity labels): (1) upregulation of apoptosis (apoptosis up); (2) cytotoxicity as measured using beta-lactamase activity as a viability reporter^27^ (cytotoxicity BLA); (3) cytotoxicity measured using SulfoRhodamine B assays that quantify cellular density based on the protein content^27^ (cytotoxicity SRB); (4) ER stress; (5) heat shock; (6) microtubule upregulation; (7) upregulation of mitochondrial disruption; (8) upregulation of oxidative stress; and (9) decrease in proliferation. We cross-referenced these nine biological activities with public Cell Painting profiles to focus on a dataset of 658 structurally unique compounds. For each of the nine biological activities, we trained Random Forest classifiers using 827 Cell Painting features to build predictive models and calculated feature importance for each Cell Painting feature. For eight out of nine biological activities (mitochondrial disruption was excluded because its models recorded AUC<0.50 and were not interpreted), the Cell Painting features most contributing to the eight models was mapped into BioMorph terms revealing interesting details about the associations between morphological features, phenotypic characteristics and cellular processes, as shown for the endpoint “ER stress” in Figure 3 for illustrative purposes. In this example, the BioMorph space terms that contain the highest percentage overlap with the Cell Painting features associated with the ER stress revealed potential secondary mechanisms of “ER stress” biological activity, such as G_2_ cell cycle arrest (level 3) and the JAK/STAT signalling pathway (level 4), both in agreement with the literature.^30,37^

**Figure 3.**
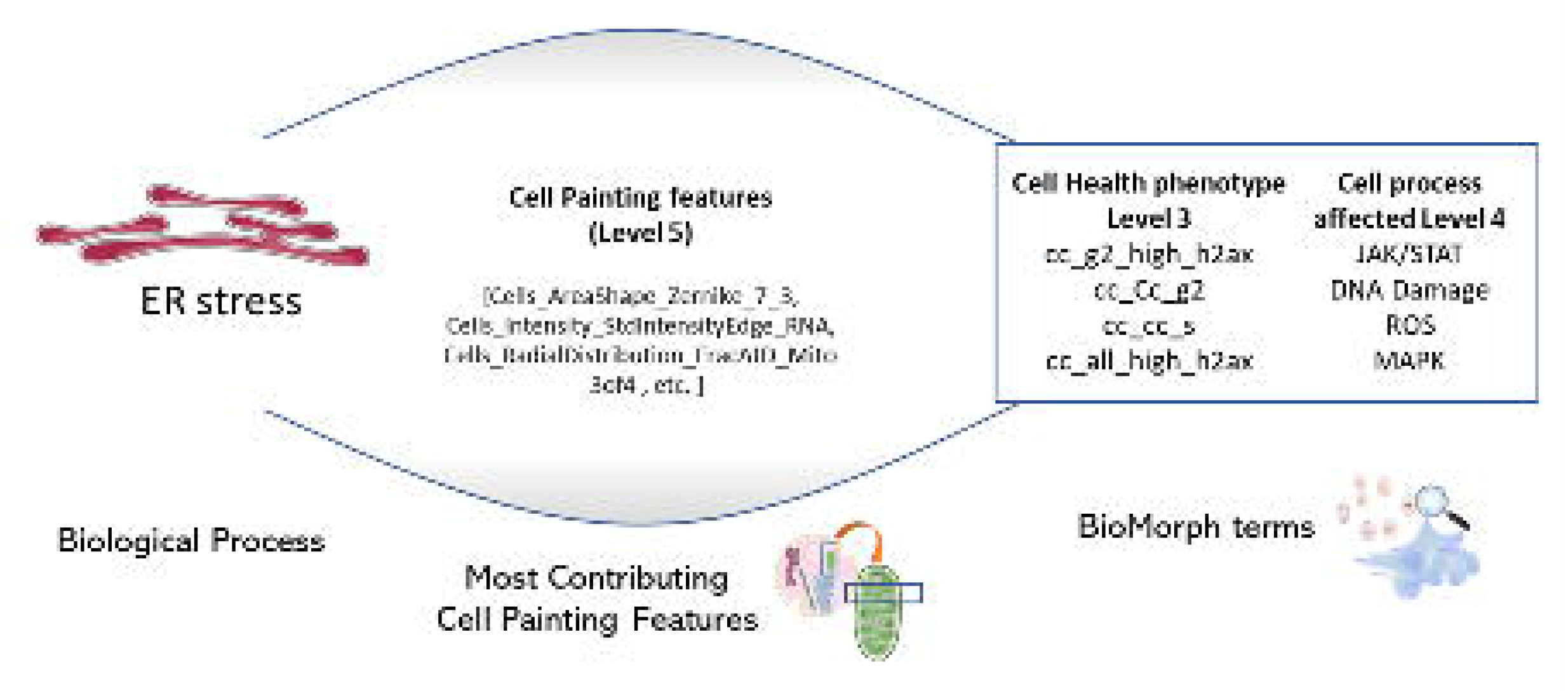
The subset of most-contributing Cell Painting features (level 5) for the model predicting ER stress and the BioMorph terms enriched from this subset (BioMorph terms that contain the highest percentage overlap with these Cell Painting features). This revealed potential secondary mechanisms of biological activity such as G_2_ cell cycle arrest the JAK/STAT signalling pathway was the most enriched Cell Health phenotype (level 3) and Cell process (level 4) respectively for ER stress.

At the level of phenotypic characteristics, the five most-contributing Cell Health phenotypes (level 3) for the eight biological processes are shown in Figure 4 (with a comprehensive analysis across various levels of BioMorph terms given in Supplementary Table S3). For the biological process of apoptosis, the most-contributing Cell Health phenotype (level 3) was the fraction of cells containing more than three γH2AX spots per cell, indicating DNA damage (Figure 4). This finding is consistent with our understanding of apoptosis as a coordinated response to DNA damage.^28^ In terms of cytotoxicity predictions, we observed that the performance of predicting results of BLA assays was improved when the BioMorph terms that incorporate Cell Health phenotypes (level 3) related to DNA damage for cells in S and G_2_ phases (Figure 4), in agreement with the well-established effect of DNA damage on cell cycle arrest. On the other hand, SRB assays measure protein content, which is affected by overall cell death, including non-apoptotic cell death, and we observed that Cell Painting features contributing to model performance here incorporated caspase-negative death Cell Health phenotypes (Figure 4). The Cell Health phenotypes (level 3) that contributed the most to the biological activities of ER stress, heat shock, and proliferation decrease were related to high γH2AX activity (based on the feature related to the fraction of G2 cells with >3 [H2Ax spots within nuclei, Figure 4), indicating DNA damage. This is consistent with previously reported observations that ER stress and heat shock cause cell cycle arrest at both G_1_/S and G_2_/M phases.^29,30,31^ For the biological activity of microtubule upregulation, the most-contributing Cell Health phenotypes (level 3) were the overall DNA damage and the fraction of caspase-negative dead cells, in agreement with their roles in cell death.^32^ Finally, for the biological activity of oxidative stress, the most-contributing Cell Health phenotype (level 3) was the average nucleus roundness, which is consistent with the significant crosstalk between DNA damage, oxidative stress, and nuclear shape alterations.^33^ Taken together, we found that the BioMorph space (level 3 Cell Health phenotypes) effectively captured biologically relevant information, allowing for a more nuanced understanding of how biological processes overall affect specific cellular processes. This is particularly advantageous compared to using Cell Painting features directly where no measurements on cell cycle phase or cell processes are made directly.

**Figure 4.**
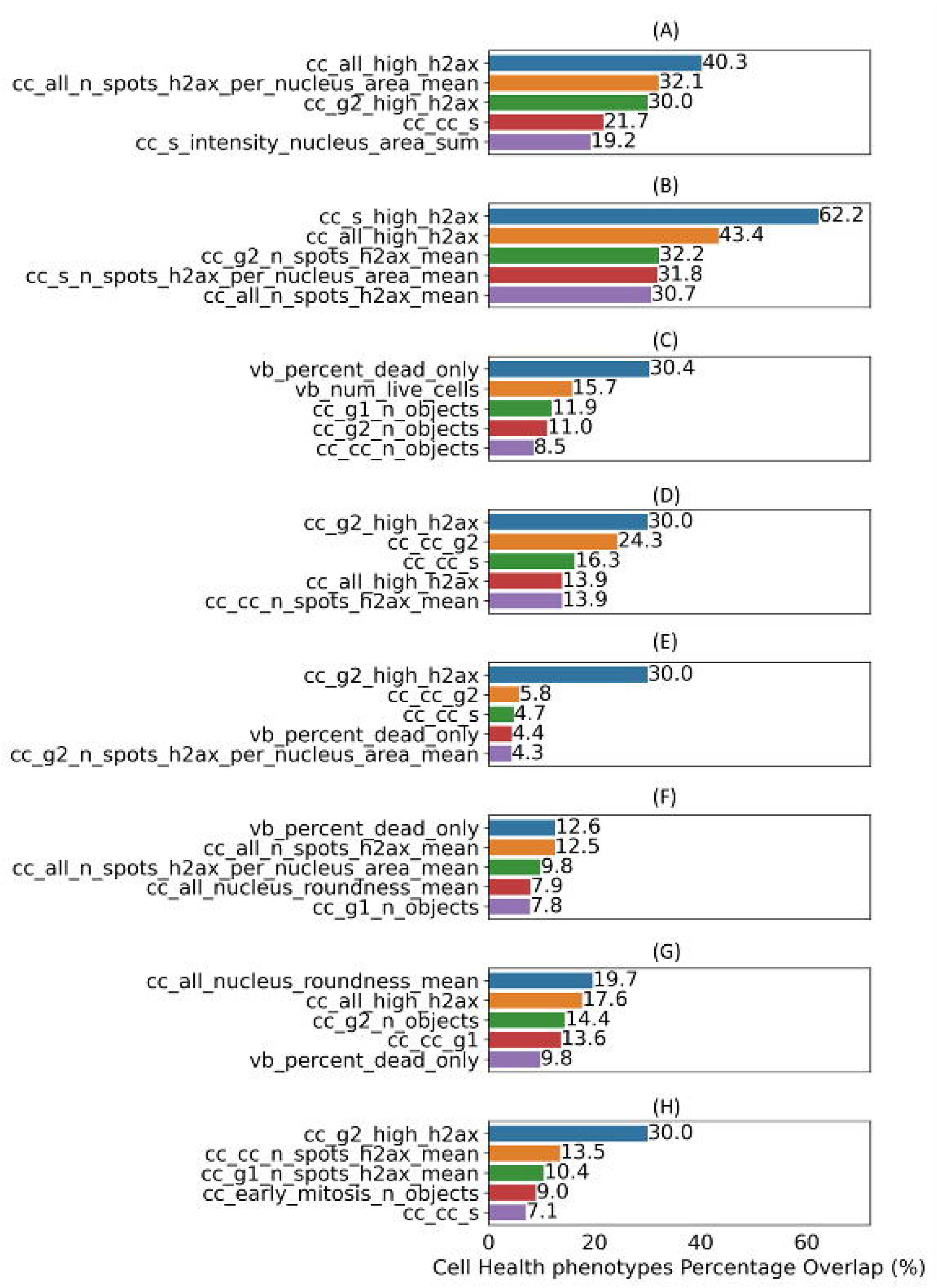
Top five specific Cell Health phenotypes (level 3) enriched by contributing Cell Painting features (as per feature importance) for each Random Forest model for eight different biological activities (a) apoptosis up, (b) cytotoxicity BLA, (c) cytotoxicity SRB, (d) ER stress, (e) heat shock, (f) microtubule upregulation, (g) oxidative stress, and (h) proliferation decrease. Models for mitochondrial disruption recorded AUC<0.50 and were not interpreted.

### Integrating information about the Cell process affected (level 4) enhances insights into mechanisms of biological activity

In addition to the information about phenotypic characteristics, the BioMorph space also includes information about specific cellular processes responsible for the alterations in cell morphology, which in turn can help to identify potential targets and biological pathways that, when modulated, could lead to desired phenotypic changes. Therefore, we examined information from affected cellular processes (level 4 of the BioMorph Space) for each of the eight biological activities. The top five enriched Cell processes associated with each of the eight biological activities are shown in Figure 5, with a comprehensive analysis across various levels of BioMorph terms given in Supplementary Table S3. For each of the eight endpoints, we found consistent agreement between the top enriched Cell processes and the existing literature. For example, in the case of apoptosis endpoint, the top three enriched processes were ROS, receptor tyrosine kinase (RTK) and mitogen-activated protein kinase (MAPK) pathways (Figure 5), which agrees with the existing literature.^34,35,36^ The JAK/STAT signalling pathway was the most enriched Cell process for ER stress (Figure 5), aligning with its role in ER stress-induced inflammation.^37^ Similarly, the most enriched processes for the other endpoints (Figure 5), *i.e.* Cytotoxicity BLA (Hippo signalling pathway), Cytotoxicity SRB (cyclosporine binding protein), heat shock response (DNA damage), oxidative stress (apoptosis and hypoxia), and proliferation (Hippo pathways), are all in agreement with expectations based on the prior knowledge.^38,39,40,41,42^ Collectively, these findings illustrate the high level of agreement between BioMorph terms and well-established biological knowledge. They also highlight how integrating information about biological processes (level 4 in BioMorph space) allows for more mechanistic interpretations and predictions.

**Figure 5.**
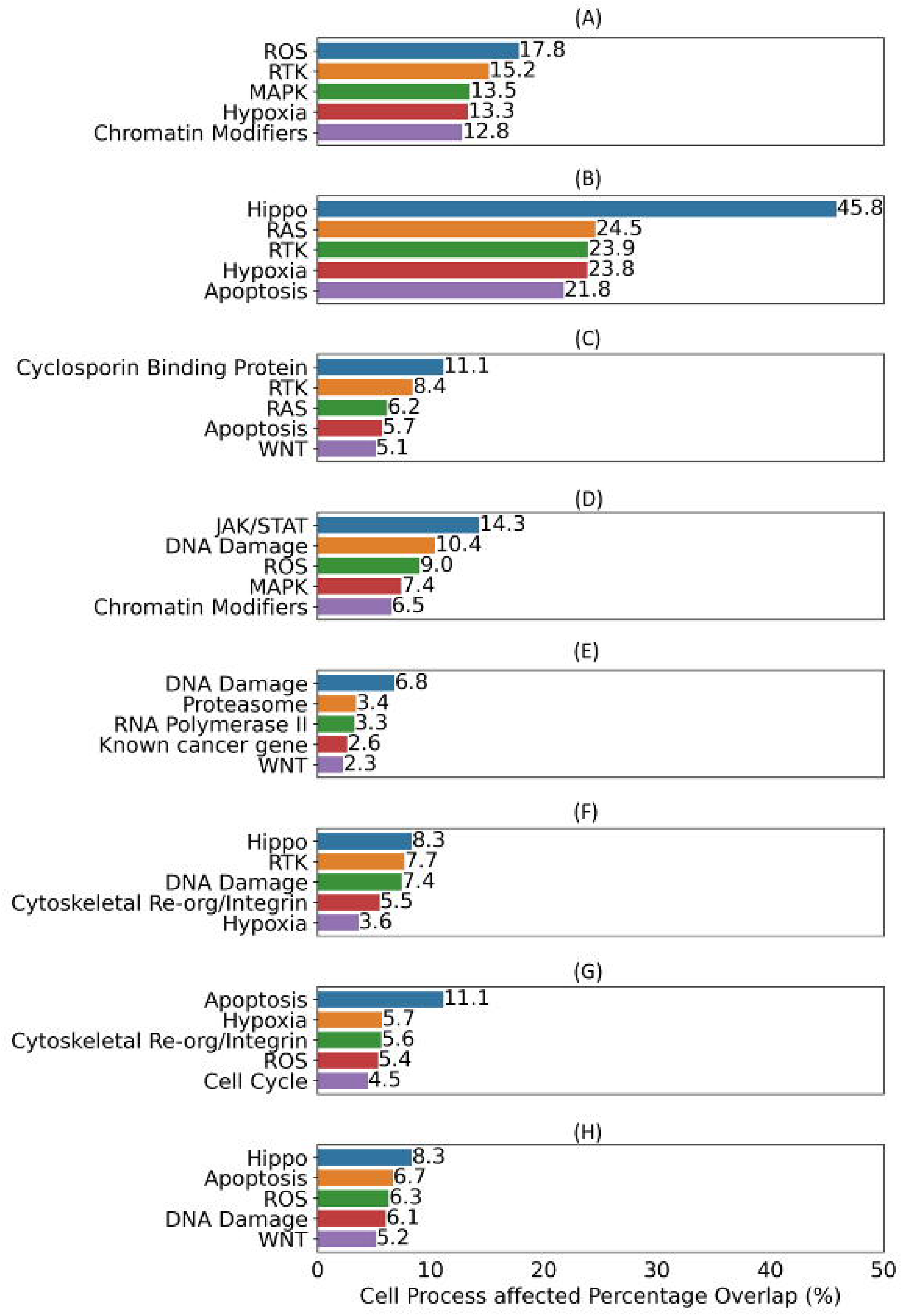
Top five Cell process affected (level 4) terms enriched by contributing Cell Painting features (as per feature importance) for each Random Forest model for eight different biological activities (a) apoptosis up, (b) cytotoxicity BLA, (c) cytotoxicity SRB, (d) ER stress, (e) heat shock, (f) microtubule upregulation, (g) oxidative stress, and (h) proliferation decrease. Models for mitochondrial disruption recorded AUC<0.50 and were not interpreted.

### BioMorph terms can be used to generate hypotheses for a compound’s mechanisms of action

We next investigated how BioMorph terms can reveal more specific mechanisms of action of a compound causing a particular biological activity. To this end, we analysed 56 predicted true positive compounds across nine biological activities and analysed the SHapley Additive exPlanations (SHAP^43^) values of Cell Painting features (a positive SHAP value for a feature indicates a positive impact on prediction, leading the model to predict toxicity in this case). These contributing Cell Painting features were mapped to the BioMorph terms, along with the two most-contributing Cell Health phenotypes (level 3 of the BioMorph) and Cell process affected (level 4 of the BioMorph). We were able to identify relationships between specific compounds and their impact on cellular health (see Table 1 for a selection of illustrative compounds discussed below; and Supplementary Table S4 for the complete set of 54 compounds analysed). For example, for melatonin, an “apoptosis up” compound, we noted that the most contributing Cell Painting features were related to BioMorph terms for DNA damage (as indicated by the presence of more than three γH2AX spots within the cells) and the fraction of cells arrested in the S phase, which is most likely due to increased ROS. In the case of melatonin, the effects on the cell cycle via ROS generation have been previously reported.^44^ In general, we observed that BioMorph space can help generate hypotheses to uncover secondary effects that might otherwise be overlooked, and examples listed in Table 1 and shown in Figure 6 speak to the granularity of the BioMorph space information. In the case of ER stressors, piromidic acid, clozapine, bisphenol A diglycidyl ether, and emetine, the top 2 most contributing Cell Health phenotypes and top 2 Cell processes affected were mostly different. This highlights that each compound may exhibit the same bioactivity (e.g. “ER stress”), but cause it by affecting different targets/pathways and having distinct MOAs. The most contributing BioMorph terms for piromidic acid are related to cell viability (such as the number of cells and roundness of living cells); whereas emetine, a protein synthesis inhibitor, was linked to the fraction of cells in the S-phase of the cell cycle, which agrees with the secondary activity in early S-phase related to inhibition of DNA replication.^45^ On the other hand, compounds linked to heat shock responses (alfadolone acetate, suxibuzone, and diflorasone) exhibited the same features and were associated with Hippo pathway-related terms, the roundness of the nucleus, and DNA damage in the S phase. This agrees with the established role of the Hippo pathway in promoting cell survival in response to various stressors^46^ while the shape of the nucleus (senescent cells can be characterized by flattened, enlarged or irregular-shape nuclei^47,48^) and vulnerability of early S-phase cells to mild genotoxic stress are common mechanisms of heat stress effects.^49,50^ We also noted similarities among the level 3 and level 4 BioMorph space terms associated with compounds that cause proliferation decrease (raclopride, nimodipine and ketanserin). These compounds are associated with hypoxia and apoptosis, suggesting that these compounds may act via increasing levels of ROS, which leads to oxidative stress.^41^ For the compounds causing an upregulation of microtubules, bifemelane was linked to BioMorph terms related to cell death as well as chromatin modifiers and DNA damage in the S phase consistent with its known role in enhancing the synthesis of cytoskeletal proteins^51^ and regulating dynamic chromosome organization.^52^ Taken together, we showcase how identifying the BioMorph terms having the greatest contribution to predicting a compound’s biological activity, we can gain insights into not only primary but secondary biological processes affected by the compounds as well. These predictions can then be used to formulate mechanistic hypotheses and inform drug discovery and development efforts.

**Figure 6.**
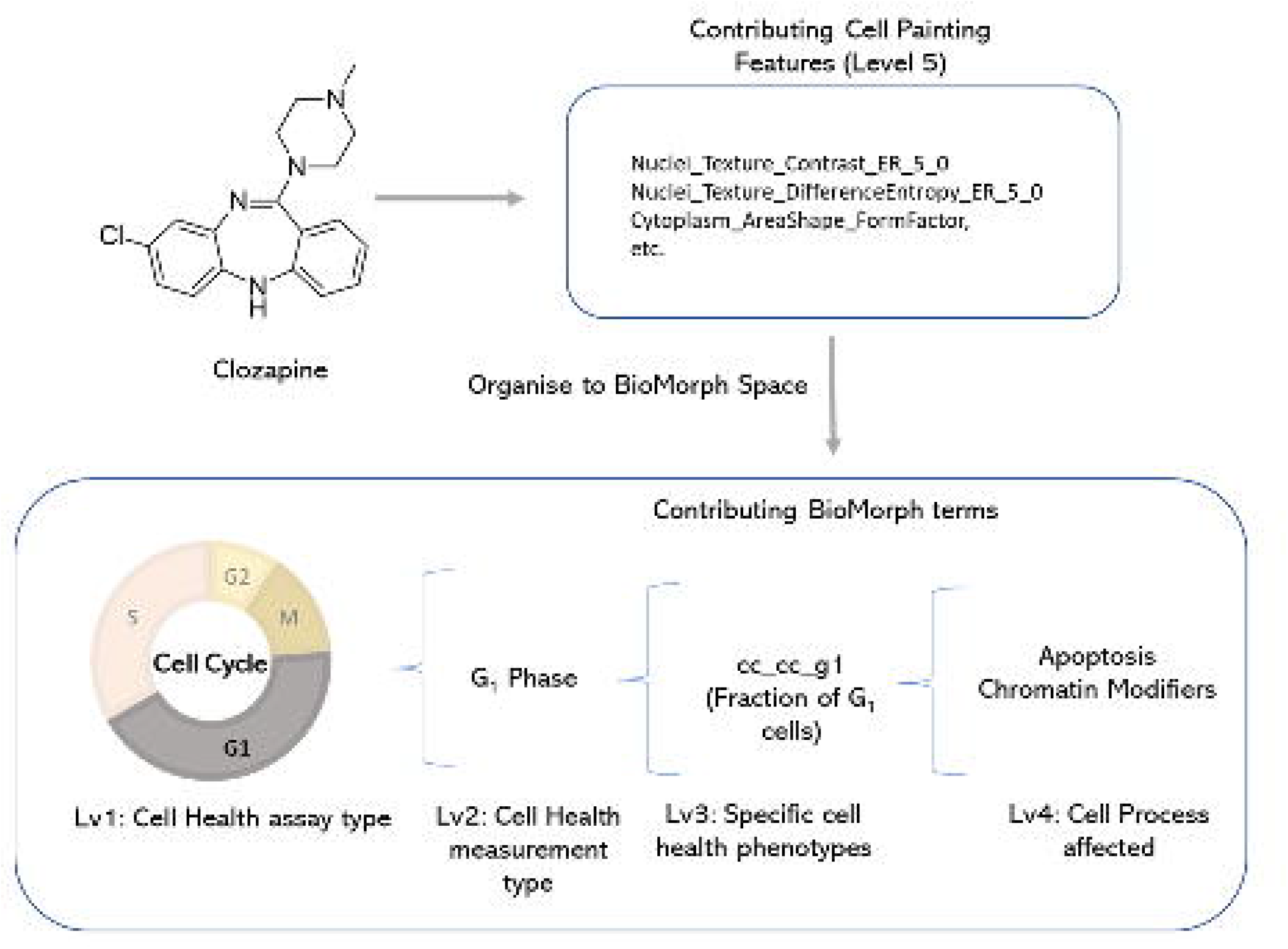
For the compound clozapine, which is an ER stressor, SHAP values indicate a list of the most-contributing Cell Painting features (Level 5) to model performance for ER stress. Organising this to BioMorph terms allows interpretation: clozapine can induce cell cycle arrest in the G_0_/G_1_ phase.

**Table 1:** Top two contributing Cell Health phenotypes (level 3) and Cell process affected (level 4) from BioMorph space for a selection of illustrative true positives predicted by the models for biological activity. See Supplementary Table S4 for the complete set of 54 compounds.

| Common name | Biological Process | Specific Cell Health phenotypes (most impacted) | Specific Cell Health phenotypes (2nd most impacted) | Cell process affected (most) | Cell process affected (2nd most) |
| --- | --- | --- | --- | --- | --- |
| Melatonin | apoptosis up | The fraction of cells containing more than 3 gH2AX spots within all cells: | Fraction of EdU positive cells (S-phase of the cell cycle) | ROS | MAPK |
| Piromidic acid | ER stress | Total number of cells | Cell Roundness | RTK | ER Stress/UPR |
| Clozapine | ER stress | Fraction of G1 cells | Fraction of caspase negative in dead cells | Chromatin Modifiers | ER Stress/UPR |
| Bisphenol A diglycidyl ether | ER stress | Fraction of G2 cells | Total number of cells | JAK/STAT | WNT |
| Emetine | ER stress | Fraction of EdU positive cells (S-phase of the cell cycle) | The fraction of cells containing more than 3 gH2AX spots within all cells | ROS | Chromatin Modifiers |
| Alfadolone acetate | heat shock | Average nucleus roundness | The fraction of >3 gH2Ax spots in S phase cells | Hippo | Chromatin Modifiers |
| Suxibuzone | heat shock | Average nucleus roundness | The fraction of >3 gH2Ax spots in S phase cells | Hippo | Chromatin Modifiers |
| Diflorasone diacetate | heat shock | Average nucleus roundness | The fraction of >3 gH2Ax spots in S phase cells | Hippo | Chromatin Modifiers |
| Bifemelane | microtubule up | Fraction of caspase negative in dead cells | EdU incorporated (average intensity per cell) in S phase cells | Hippo | Chromatin Modifiers |
| Raclopride | proliferation decrease | Fraction of caspase negative in dead cells | Width/Length | Hypoxia | Apoptosis |
| Nimodipine | proliferatio<br>n decrease | Fraction of<br>caspase negative<br>in dead cells | Width/Length | Hypoxia | Apoptosis |
| Ketanserin | proliferatio<br>n decrease | Number of G1<br>cells | Fraction of caspase<br>negative in dead cells | Hypoxia | Apoptosis |

### Limitations of mapping Cell Painting into BioMorph terms

This proof-of-concept study demonstrates the potential benefits of mapping Cell Painting features into BioMorph terms to address a serious challenge for the field of image-based profiling: making sense of complex combinations of image-based features that are not readily interpretable. We find that BioMorph does provide a more interpretable and biologically relevant representation of data. However, there are several limitations relevant to this iteration of BioMorph space. BioMorph space was built using robust but limited data; therefore, using larger datasets of CRISPR perturbations and Cell Painting / Cell Health datasets would improve the organization of BioMorph space. Additionally, the associations between Cell Painting features and BioMorph terms are not absolute; these would need to be updated if alternative feature extraction strategies are used (e.g., updated versions from CellProfiler or deep learning-based feature extraction such as in the JUMP-Cell Painting dataset^53^), and we advise caution against using these groupings directly if the datasets differ from the current study. Finally, we evaluated our BioMorph space for nine broad biological activities; generalization to other cellular mechanisms and biological processes would require assays that are focused on other readouts, such as those related to particular types of toxicity, or tailored to particular cell types like neurons or cardiomyocytes. Despite these limitations, the study introduces an algorithm to map Cell Painting features into BioMorph terms and explores the application of this new BioMorph space in interpreting predictive models, generating hypotheses for small molecule biological activity, MOA, and toxicity.

## SIGNIFICANCE

In this work, we demonstrated a strategy to map Cell Painting features into BioMorph terms to enable a better understanding of the relationships between compound-induced cellular perturbations and nine different biological activities. We could correctly identify potential secondary mechanisms of biological activities such as ER stress and cell cycle arrest at the G_2_ phase^30^ as well as mechanisms of action of dual-function compounds such as emetine, which is a well-known protein synthesis inhibitor, but also acts at an early S-phase to inhibit DNA replication^45^. These are biological effects that can often be overlooked; however, the BioMorph space allows for a more comprehensive understanding of these mechanisms, for uncovering hidden relationships and generating new hypotheses by connecting them to specific phenotypes and cellular processes.

Mapping Cell Painting features into BioMorph terms offers several advantages over using Cell Painting features directly. First, we improved interpretability by using a more biologically interpretable feature space; we identified relationships between compound mechanisms of action and their impact on cell morphology. For example, the use of BioMorph space identified relevant pathways such as the JAK/STAT signalling pathway’s prominence in ER stress.^37^ These insights are not possible with Cell Painting features alone, which have no information on biological pathways. Second, we could pinpoint the specific cell processes and stages of the cell cycle affected by a compound, a task not possible with the Cell Painting features, which do not contain direct information on which cell cycle stage is impacted. Finally, we could facilitate hypothesis generation by identifying the BioMorph terms that contribute most significantly to compound activity. These targeted hypotheses can guide the future validation of compounds. Taken together, the BioMorph space represents a more integrative and comprehensive method for analysing cellular MOA and can enable the development of more effective strategies for identifying and mitigating toxic effects.

## METHODS

### Cell Painting Dataset for CRISPR Perturbations

We used the Cell Painting pilot dataset of (CRISPR) knockout perturbations from the Broad Institute.^6**Error! Bookmark not defined.**^ Here, the authors used a Cell Painting assay for three different cell lines (A549, ES2, and HCC44) and each cell line used 357 perturbations representing 119 clustered regularly interspersed short palindromic repeats (CRISPR) knockout perturbations (further details in Supplementary Table S5). They further generated median consensus signatures for each of the 357 perturbations. This led to a dataset of 949 morphology features (and metadata annotations) for 357 consensus profiles (119 CRISPR perturbations × 3 cell lines). Among these, only 827 Cell Painting features were in intersection with the Cell Painting dataset for compound perturbations (described below) used in this proof-of-concept study. The Cell Painting dataset for CRISPR Perturbations is released publicly at https://zenodo.org/record/8147310.

### Cell Health assays for CRISPR Perturbations

We used the Cell Health assay developed by the Broad Institute containing 70 specific Cell Health phenotypes.^6^ The authors used seven reagents in two Cell Health panels to stain cells for the same 119 CRISPR perturbations for three different cell lines (A549, ES2, and HCC44). We used median consensus signatures for the 357 consensus profiles (119 CRISPR perturbations × 3 cell lines) as above. This dataset is released publicly at https://zenodo.org/record/8147310.

### Cell Painting Dataset for Compound Perturbations

The Cell Painting assay used in this proof-of-concept study, from the Broad Institute, contains cellular morphological profiles of more than 30,000 small molecule perturbations.^22^ The morphological profiles in this dataset are composed of a wide range of feature measurements (share, area, size, correlation, texture etc). The authors in this study normalized morphological features to compensate for variations across plates and further excluded features having a zero median absolute deviation (MAD) for all reference cells in any plate. Following the procedure from Lapins et al^54^, we subtracted the average feature value of the neutral DMSO control from the compound perturbation average feature value on a plate-by-plate basis. We standardised the InChI using RDKit^57^ and for each compound and drug combination, we calculated a median feature value. Where the same compound was replicated for different doses, we used the median feature value across all doses that were within one standard deviation of the mean dose. Finally, we obtained 1,783 median Cell Painting features for 30,404 unique compounds. Among these, only 827 Cell Painting features were common with the dataset for CRISPR Perturbations which were used in this proof-of-concept study.

### Biological activity from ToxCast assay with Cell Painting annotations

Toxicity and biological activity-related data were collected from 56 cytotoxicity and cell stress response assays from 56 ToxCast^26,55^ for nine broad biological processes(for the mapping between 56 ToxCast assays and 9 biological processes see Judson et al^56^): apoptosis up, cytotoxicity BLA, cytotoxicity SRB, ER stress, heat shock, microtubule upregulation, mitochondrial disruption up, oxidative stress up and proliferation decrease.^56^ Compound SMILES were converted to standardised InChI using RDKit.^57^ To generate consensus endpoint labels, the presence of positive activity (toxicity) in at least one assay related to the biological activity was considered sufficient to mark the compound active in the consensus endpoint. Thus, consensus endpoints for each of the 9 biological activities were generated from the 56 ToxCast assays. We calculated the intersection of the Cell Painting profiles for compound perturbations (above) and 9 biological activity (ToxCast) assays using the standardised InChI. Cell Painting features were standardised by removing the mean and scaling to unit variance. This resulted in a complete dataset of 658 structurally unique compounds with 827 Cell Painting features and 9 biological activity consensus hit calls that were used in this proof-of-concept study. The dataset, referred to as containing biological activities in this study, is publicly released in https://zenodo.org/record/8147310.

### Mapping Cell Painting terms into BioMorph space

The overlap of Cell Painting and Cell Health assay for gene perturbations from Way et al^6^ contained 827 Cell Painting features (that were also present in the Cell Painting experiments on compound perturbations from Bray et al^4^) and 70 continuous Cell Health endpoints (e.g., the number of late polynuclear cells, which measures the shape in a cell cycle assay) for 354 consensus profiles (118 CRISPR perturbations × 3 cell lines, the empty well was removed). As shown in Figure 2 step A, for feature selection, we used an all-relevant feature selection method, Borutapy^58^ (implemented using the Python package Boruta^59^) with a Random Forest Classifier estimator of maximum depth 5 and the number of estimators determined automatically based on the size of the dataset using ‘auto’. Using Borutapy, we detected a subset of Cell Painting features that contain information for each of the 70 Cell Health regression labels. Further, we trained a baseline Linear Regression model as implemented in scikit-learn^60^ (Figure 2 step B) with an 80-20 random train-test split to predict which subsets of Cell Paintings features are relatively better predictors of Cell Health phenotype. 37 of the 70 Cell Health models (with R^2^>0.25) were selected for further analysis. This results in 34 subsets of Cell Painting features (one set for each of the 34 Cell Health labels). Next, for each of the 34 Cell Health labels and the 354 consensus profiles, we separated subsets of Cell Painting data for the negative control CRISPR Perturbation (which consisted of 30 datapoints of LacZ, Luc, Chr2 CRISPR perturbations) and other CRISPR perturbations affecting various known cell processes (such as chromatin modifiers, ER Stress/UPR, metabolism etc). For each of these pairs (negative control and CRISPR perturbations), we used Borutapy^58^ (Figure 2 step C) to detect a further subset from the subset of Cell Painting features which contained a signal on whether the datapoint is a negative control or the CRISPR Perturbation. We train a baseline Random Forest Classifier (Figure 2 step D), as implemented in scikit-learn^60^, with an 80-20 random train test split to predict which sets of selected Cell Painting features perform relatively better at differentiating negative controls from the CRISPR perturbation (MCC>0.50). This led to 412 subsets of informative Cell Painting features which are then indicators of 412 BioMorph terms. We used a χ2 test to determine the BioMorph term p-value for each of the 412 combinations (Figure 2 step E) from standard scaled subsets of Cell Painting features.

Further using this mapping, any dataset with Cell Painting features can be mapped into BioMorph terms. The dataset of biological activities with 827 Cell Painting features were grouped into these 412 combinations and their BioMorph term p-value was calculated. We then standardised these BioMorph terms using a standard scalar (as implemented in scikit learn), and only columns with non-infinite continuous p-values were retained (with other columns dropped). This resulted in 398 BioMorph terms for the biological activity dataset. The dataset is now publicly released in https://zenodo.org/record/8147310.

### Comparing models using Cell Painting and BioMorph terms as features

To ensure that the BioMorph terms contain all information from the original Cell Painting readouts, we compared models using only 827 Cell Painting features and models using the 398 BioMorph terms directly as features (although there were 412 terms defined, only 398 terms out of these were non-infinite and continuous and used for modelling). For each of the 9 biological activities, we used 5 times repeated 4-fold nested cross-validation and a Random Forest Classifier (as implemented in scikit-learn^60^). First, the data was split into four folds using a stratified split on biological activity labels where 25% of the data was reserved for the test set and 75% remaining used for training. Using this training data, we trained two models, one using the 827 Cell Painting features, and the other using 398 BioMorph terms (p-values from subsets of Cell Painting features; although there were 412 terms defined, only 398 terms out of these were non-infinite and continuous and used for modelling). We optimised these models using a 5-fold cross-validation with stratified splits and a random halving search algorithm (with hyperparameter space given in Supplementary Table S6 and as implemented in scikit-learn^60^). The optimised model was fit on the entire training data and cross-validation predictions are used to determine the optimal threshold using the J statistic value. We then used this threshold to determine the predictions for the test set predictions. A single loop of nested cross-validation results in 4 test sets, which are repeated 5 times thus giving 20 individual test set predictions.

### Model training with Cell Painting features

To evaluate the use of BioMorph space, we now used a fixed held-out test set. For each of the 9 biological activities, we used a stratified split on biological activity labels such that 75% of the data was used in cross-validation training and 25% as held-out test data. We trained Random Forest classifiers (as implemented in scikit-learn^60^) using 827 Cell Painting features and a random halving search algorithm (as implemented in scikit-learn^60^) to optimise the hyperparameters (with the hyperparameter space given in Supplementary Table S6). Similar to above, the optimised model was fit on the entire training data and cross-validation predictions are used to determine the optimal threshold using the J statistic value that considers both true and false positive rates. This optimal threshold is then used on the predicted probabilities of the held-out test data to obtain the final held-out test data predictions.

### Feature importance and interpretation in BioMorph terms

First, we next used feature importance from the Random Forest classifier (as implemented in scikit-learn^60^) to determine the features that contributed the most to model importance. This gave us important features per biological activity (at an endpoint/biological activity level). Second, we evaluated SHAP^43^ values (as implemented in the shap^61^ python package) for each compound predicted as true positive in the held-out test set. We used true positives only, as these are the predictions for which the feature importance value (from SHAP) is valid. This gave us the important features per toxic compounds in the held-out test set for each biological activity (at a compound level). We then selected the Cell Painting features (from model importance values at the endpoint level or SHAP values at a compound level) that were greater than two standard deviations of all features as the most important or contributing features. These features were mapped into the BioMorph space by determining if the features related to the individual levels of the BioMorph term were present among the important features selected above. At the level of Cell process affected (level 4), the percentage enrichment was determined as the percentage of Cell Painting features that were present among the defined subset of Cell Painting features (level 5). For an overall enrichment value (used for Figure 3 and Figure 4) for each specific Cell Health phenotypes term (level 4) or Cell process affected (level 5), we used the mean of enrichment of all BioMorph terms where the corresponding level 3 or level 4 term appeared. For detailed enrichment analysis, we determined enrichment of the level (lv_X_) to be the percentage of the immediate lower level (lv_X-1_) with enrichment ≥ 10% progressively from specific Cell Health phenotypes (level 3) to Cell Health assay type (level 1). This is released per biological activity in Supplementary Table S3.

### Evaluation Metrics

To evaluate models in this proof-of-concept study we used Balanced Accuracy which considers both sensitivity and specificity, the Area Under Curve-Receiver Operating Characteristic (AUC-ROC) and Mathew’s correlation constant (MCC) as implemented in scikit-learn^60^.

### Statistics and Reproducibility

We have released the datasets used in this proof-of-concept study which are publicly available at https://zenodo.org/record/8147310. We released the Python code for the models which are publicly available at https://github.com/srijitseal/BioMorph_Space.

## ACKNOWLEDGEMENTS

S.S. acknowledges funding from the Cambridge Commonwealth, European and International Trust, Boak Student Support Fund (Clare Hall), Jawaharlal Nehru Memorial Fund, Allen, Meek and Read Fund, Trinity Henry Barlow (Trinity College) and Cambridge Centre for Data-Driven Discovery (C2D3) and Accelerate Programme for Scientific Discovery. AEC acknowledges funding from the National Institutes of Health (R35 GM122547). OS acknowledges funding from the Swedish Research Council (grants 2020-03731 and 2020-01865), FORMAS (grant 2022-00940), Swedish Cancer Foundation (22 2412 Pj 03 H), and Horizon Europe grant agreement #101057014 (PARC) and #101057442 (REMEDI4ALL).

## ASSOCIATED CONTENT

### Supplemental Information

The Supporting Information is available. Supporting Information (PDF). We released the Python code for our models which are publicly available at https://github.com/srijitseal/BioMorph_Space/ and all datasets at https://zenodo.org/record/8147310.

Supplementary Table S1: The groups of Cell Painting features for each of the 412 BioMorph terms along with a description of the specific phenotype used in this study.

Supplementary Table S2: Mean evaluation metrics from repeated nested cross-validation for both Random Forest classifiers using (a) 827 Cell Painting features and (b) 398 BioMorph terms for 9 biological activity endpoints used in this study.

Supplementary Table S3: The detailed analysis of each of the eight biological activities and their enrichment at various levels of BioMorph terms, progressively from specific Cell Health phenotypes (level 3) to Cell Health assay type (level 1).

Supplementary Table S4: The BioMorph terms that positively impact all 56 compounds predicted true positives for 9 biological activity endpoints used in this study.

Supplementary Table S5: The Cell process affected by individual CRISPR knockout perturbations from the Cell Health and Cell Painting dataset used in this study.

Supplementary Table S6: Distribution of hyperparameters used to optimise Random Forest models used in this study.

## AUTHOR INFORMATION

### Corresponding Authors

Prof Andreas Bender, Prof Ola Spjuth

All authors have approved the final version of the manuscript.

### Contributions

S.S. designed and performed data analysis, constructed the BioMorph space organisation, and implemented, and trained the models. S.S, J.C.P and O.S analysed the biological interpretation of morphological features. S.S. and J.C.P. contributed to analysing the results of models. S.S. wrote the manuscript with extensive discussions with O.S., J.C.P and A.E.C.. A.B. and O.S. supervised the project. All the authors (S.S, J.C.P, A.E.C, O.S and A.B) reviewed, edited, and contributed to discussions on the manuscript.

### Conflicts of Interest

The authors declare no conflict of interest.

## Notes

https://zenodo.org/record/8147310

https://github.com/srijitseal/BioMorph_Space

### Competing Interest Statement

The authors have declared no competing interest.

