## Supplementary Figures for "From Pixels to Phenotypes: Integrating Image-Based Profiling with Cell Health Data Improves Interpretability"

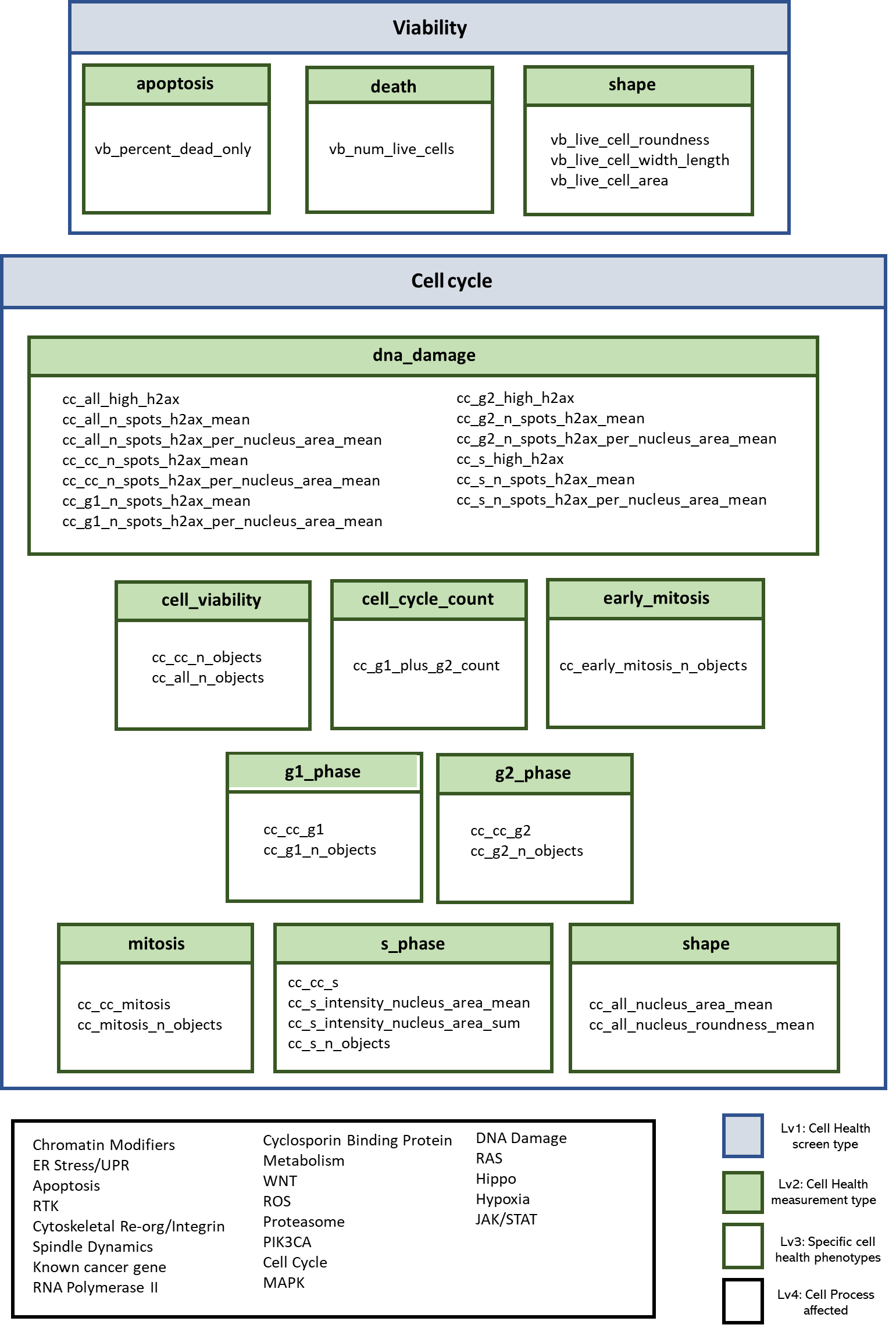

Figure S1: BioMorph terms’ hierarchy with relationships for (a) Level 1: Cell Health assay type, (b) Level 2: Cell Health measurement type, (c) Level 3: Specific Cell Health phenotypes, (d) Level 4: Cell process affected (representative of morphological characteristics)

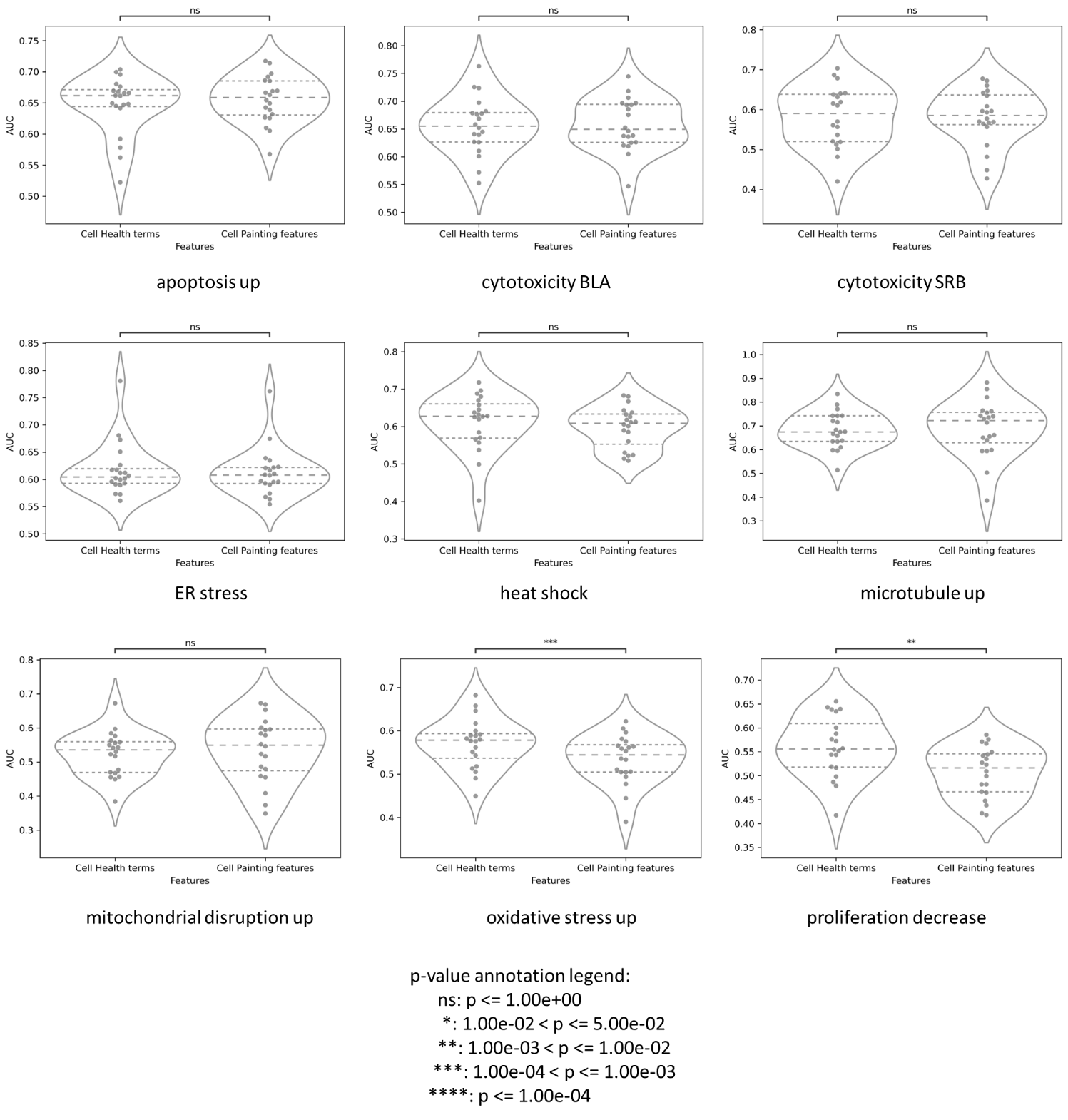

Figure S2: Distribution AUC for both models, namely, Cell Painting features and BioMorph terms as features when predicting 9 different broad biological activities in 5 times repeated 4-fold nested cross-validation.
